# Structural characterizations of a unusual peptide CORO1C-47aa encoded by circRNA hsa-circ-0000437 associated with endometrial cancer

**DOI:** 10.1101/2022.12.19.521054

**Authors:** Sima Biswas, Gaurav Kumar Bhagat, Angshuman Bagchi

## Abstract

Circular RNAs, which have covalently closed ends, is a special type of non-coding RNAs. Recent studies reveal that they are associated with various biochemical pathways. One such involvements of circular RNAs is in the onset of different types of cancers. Though the circular RNAs are known as non-coding RNAs, however, some of them are found to possess the capacities to code for proteins. One such circular RNA is hsa-circ-0000437 which is known to code for a short peptide referred to as CORO1C-47aa. The peptide has anti-angiogenic activity and are associated with the prevention of endometrial cancer. However, till date only the amino acid sequence of the peptide is known and no structural details of the peptide are available. Therefore, in this work, our aim was to predict how the peptide would fold and what could be its possible ligand binding sites. We used computational tools to determine the structure of the peptide. The structure was refined by molecular dynamics simulations. The possible binding sites and the binding partners of the peptide were also predicted. From this structure function analysis study, we tried to elucidate the possible mechanism of the involvements of the peptide in the onset of endometrial cancer. This is the first report on the structural characterization of the peptide. This study may therefore be useful in determining the structures of new drug candidates for the treatment of endometrial cancer.

## 1. Introduction

Circular RNAs are covalently closed RNA molecules which belong to the class of noncoding RNAs. They are obtained by the method of backsplicing [1–5]. Initially, they were considered to be transcriptional junks. However, with the advent of the Next Generation Sequencing (NGS) technologies as well as advanced Bioinformatics algorithms a huge repertoire of circular RNA molecules have been discovered and furthermore, the tissue specificity and functionalities of these circular RNAs have started been being identified [6,7]. The fundamental classifications of the circular RNAs include the exonic, intronic or exon-intronic. The circular RNAs are stable molecules and are resistant to degradation by RNAses. Recent studies have revealed that the circular RNA molecules take part in numerous cellular activities like, Gene Expressions, Sponging of miRNAs, and Binding to proteins for the regulatory purposes to name a few [1–10]. As a matter of surprise, some of the circular RNAs have been found to be able to code for peptides/proteins having a variety of functionalities [11–15]. In this work, we tried to characterize the unusual peptide encoded by the circular RNA, circ-0000437.

The circ-0000437 is known to be associated with the onset of endometrial cancer and it is extracted from the human endometrial cancer cells [16]. Surprisingly, the apparently non-coding circularRNA is found to be able to code for a short peptide using a 144 nucleotide long ORF referred to as sORF. The encoded peptide, CORO1C-47aa, contains 47 amino acid residues. CORO1C-47aa, when over-expressed, is found to have profound influence on endothelial network formations. The peptide is also known to a) inhibit cellular angiogenesis, b) suppress the proliferation, migrations and differentiation of endothelial cells. In short, CORO1C-47aa may act as an anti-angiogenic factor for endometrial cancer [16]. It binds competitively with Aryl hydrocarbon Receptor Nuclear Translocator (ARNT). ARNT interacts with Transforming Acidic Coiled-Coil 3 (TACC3) protein and this interaction is necessary for tumor angiogenesis to promote the onset of endometrial cancer. The competitive binding of CORO1C-47aa with ARNT would lead to anti-angiogenicity and prevention of endometrial cancer [16].

Considering the profound importance of the unusual peptide CORO1C-47aa as an anti-angiogenic factor for endometrial cancer, we tried to decipher the structure-function relationship of the peptide at the molecular level. We generated the three dimensional structure of the peptide from its amino acid sequence using fold recognition method. After validations of the structure of the peptide, we performed molecular dynamics (MD) simulations on the peptide to monitor its cellular behaviors at the atomic level by measuring the Root Mean Squared Deviation (RMSD), Root Mean Squared Fluctuation (RMSF) and Radius of gyration (Rg). The changes in the secondary structural elements during the course of the MD simulations were also analyzed. The plausible ligand binding sites of the peptide were predicted. Till date, there are no reports regarding the structure-function relationship of this unusual but highly important peptide. Furthermore, only the amino acid sequence of the unusual peptide is known. We for the first time, tried to predict and analyze the different structural features of this peptide obtained from the circular RNA, circ-0000437. Our work is the first of its kind which would be able to shed light to predict the molecular mechanism of the onset of endometrial cancer as well as its preventive measures as the peptide is known to be an anti-angiogenic factor in endometrial cancer.

## 2. Materials and Methods

### 2.1. Sequence analysis of the peptide and building of model

The peptide CORO1C-47aa has the following 47 amino acid residues as obtained from the circular RNA hsa-circ-0000437 [16]. The obtained peptide sequence is as follows:

> MNPRTSTTSTHSAARSLREGWVTCPRGDLMLTNVRLPEKYAGTNCSS

We used the amino acid sequence information to search it against the Protein Data Bank (https://www.rcsb.org/) [17] to obtain similar three dimensional structures using the tool BLAST (https://blast.ncbi.nlm.nih.gov/Blast.cgi) [18]. This is performed to determine a template structure for the peptide in order to generate the three dimensional structural model of the peptide by the method of homology modelling. However, no suitable structures to be used as templates were found in the PDB. Therefore, we used the I-TASSER tool (https://zhanggroup.org/I-TASSER/) [19] to generate the modeled structure of the peptide. The three dimensional structure of the peptide was generated by using the method of multiple threading. The final model was generated by the method of fragment assembly [19].

The model of the peptide generated by the method was further refined by energy minimization to remove the steric hindrances in the modeled structure. We used Discovery Studio 2.5 (DS 2.5) for the purpose of structure refinement with the help of the Smart minimizer algorithm - a combination of steepest decent and conjugate gradient protocols. After an iteration of 5798 steps, the necessary RMS gradient of the energy derivative was obtained which was 0.001 kcal/mole. The stereo-chemical properties of the peptide were determined by SAVES6.0 server (https://saves.mbi.ucla.edu). We used the tool PROCHECK (https://servicesn.mbi.ucla.edu/PROCHECK) [20] to draw the Ramachandran plots [21] of the peptide. None of the amino acid residues of the modeled peptide were present in the disallowed regions of the Ramachandran plot. This would indicate a good model quality.

Some other physico-chemical parameters of the peptide were analyzed by considering the sequence information as present in the distribution of amino acid residues in the peptide with the help of the tool Protparam [22], ProtScale (https://web.expasy.org/protscale/) and Qmean (https://swissmodel.expasy.org/qmean/).

### 2.2. Molecular dynamics simulation study of the peptide

Molecular dynamics simulations are employed to study how a biomolecule might behave inside a virtually created cell. In order to study such behaviors we used Gromacs 5.1.5 (GROningenMAchine for Chemical Simulations) tool [23] to run MD simulations using the built model of the peptide. The atomic charges of the amino acid residues of the peptide were typed using the all atom CHARMm27 force field [24]. After that the peptide was placed in a cubic box maintaining appropriate distance from the edges of the cubic box and solvated with requisite number of simple point charge (SPC) water molecules [25].

The system was made neutral by adding necessary counter ions Na^+^ & Cl^-^. The entire system of the peptide in the solvation box was subjected to energy minimizations with the help of steepest descent algorithm. This step was performed to remove unwanted steric clashes from the built peptide model. Then the system was equilibrated in two stages each having a time step of 100ps. The first equilibration step is called the NVT, where the number of particles (N), volume (V) and temperature (T) of the entire system were kept constant. The second step, referred to as the NPT, keeps N, the pressure of the system (P) and T to be constant. The utility of the equilibration step is to make the system free from any unwanted external influence. Particle Mesh Ewald (PME) protocol was used to measure the long range electrostatic interactions of the system. The entire simulation was performed maintaining the pressure at 1 bar and temperature to the physiological temperature of 310K. The final production MD run was continued for 250ns, after the NPT step, in triplicate. This was done in order to get a comprehensive and consensus result.

The continuities of the MD simulation runs were monitored by measuring the RMSD of the backbone atoms of the peptide with the passage of time. The atomic level fluctuations of the amino acid residues of the peptide were measured by the analysis of the RMSFs during the course of MD simulation runs. We further measured how the peptide structure would deviate from its initial state with the help of Rg. In continuation of the previous analyses, we also measured the changes in the relative dispositions of the secondary structural elements of the peptide during the course of MD simulations with the tool DSSP [26].

### 2.3 Binding site analysis of the peptide

The peptide is known to interact with the protein ARNT as mentioned before. However, in order to determine the information regarding any other binding partner of the peptide, we used the following tools, viz., COACH, COFACTOR, TM-SITE and S-SITE (https://zhanggroup.org/COACH/) [27].

## 3. Results

### 3.1. Structural details of the peptide

The peptide is an unusual one as it is encoded by a so called non-coding circular RNA hsa-circ-0000437. Even after rigorous searching of the structural database PDB to find suitable template structures, we were unable to find any proper template to build the three dimensional structure of the peptide by the technique of homology modeling. In our quest to search for suitable templates, we used the Qmean server as well. The score received from Qmean server gives an idea regarding the presence of suitable structural homologs in PDB. We retrieved a small score of 0.41 ± 0.12 which would indicate that no suitable homologous structures are available in PDB

Therefore, the three dimensional structure of the peptide was generated by threading method with the help of the tool I-TASSER. The stereo-chemical qualities of the modeled peptide were detected by SAVES 6.0 server. There were no amino acid residues in the disallowed regions of the Ramachandran plot (Supplementary figures S1.A & S1.B). This would ensure a good quality of the built model. The modeled structure of the peptide is presented in Figure1.

The major secondary structural element of this peptide is helices spanning the amino acid residues 10-17 and 38-40. Two small patches of β-sheets are also present covering the amino acid residues 23-24 and 29-30. Coils and turns join the helices and sheets together. The helices are present both at the N and C termini of the peptide. The intermediate part of the peptide was found to consist of ß-sheets. The percentages of the secondary structural elements in the peptide are as follows:

Helix: 23.40%
Sheet: 8.51%,

We tried to analyze the core of the peptide to learn its structural integrity. For that we calculated the percentage buriedness and hydropathy profile of the peptide with the help of Protscale (Supplementary figures S2.A & S2.B). The peptide’s hydropathy score was positive after the 26^th^ amino acid and remained positive uptill the 28^th^ amino acid residue and from 30^th^ to 34^th^ amino acid residue as shown in S2.A. Overall, the peptide is mostly hydrophilic with some isolated patches of hydrophobic regions. The percentage buriedness score of the amino acids is presented in S2.B. These scores would signify that the buriedness of the residues was low in general. However, slighly higher scores were observed from the 9^th^ to the 20^th^ amino acid residues. This part of the peptide would be more accessible in comparison to the other parts.

Supplementary Table1 shows the results of some important physico-chemical parameters of the peptide as obtained from the ProtParam tool. The aliphatic index of the peptide was calculated as 51.91. This relatively high value of the aliphatic index would further signify the thermal stability of the peptide. The instability index of the peptide was 28.83 which would indicate that the peptide is stable. Grand average of hydropathicity (GRAVY) value was - 0.653. The negative GRAVY value of the peptide would indicate that the peptide is hydrophilic in nature.

### 3.2. Analysis of the structural fluidity of the peptide

At the end of each of the MD simulation runs, the trajectory files were analyzed to check the structural integrity of the peptide. Each of the MD simulation run was conducted for 250 ns in triplicate. After each of the the simulation runs, RMSD, RMSF, Rg, and secondary structure profiles of the peptide were analyzed. From the RMSD curve, we observed that the structure of the peptide would gain stability after 120 ns and remained stable till the end of the simulation runs (Fig 2A).

**Figure1:**
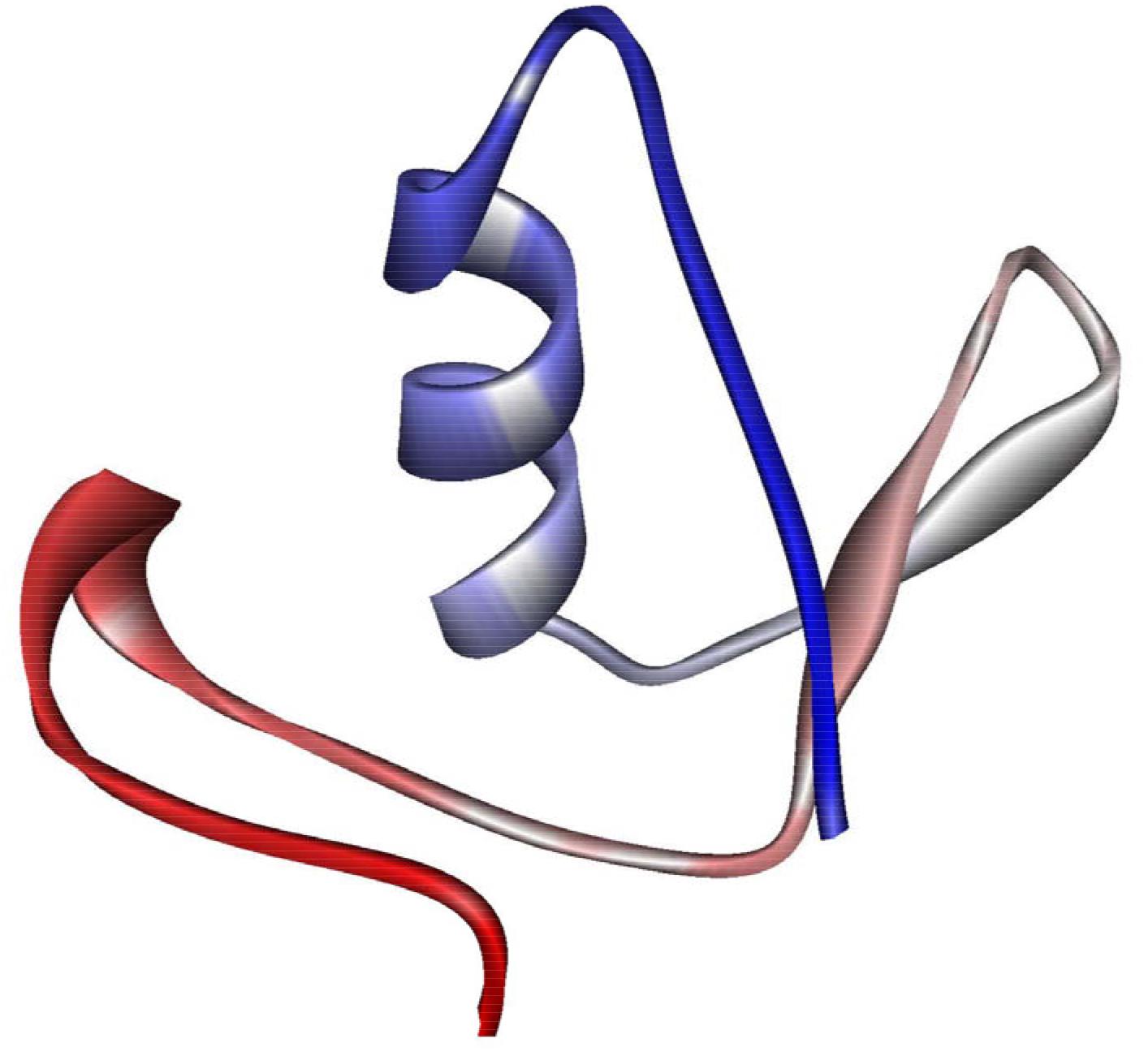
The 3D structure of the modeled protein. Colour code: Helix: Blue Red: Sheets.

**Figure2:**
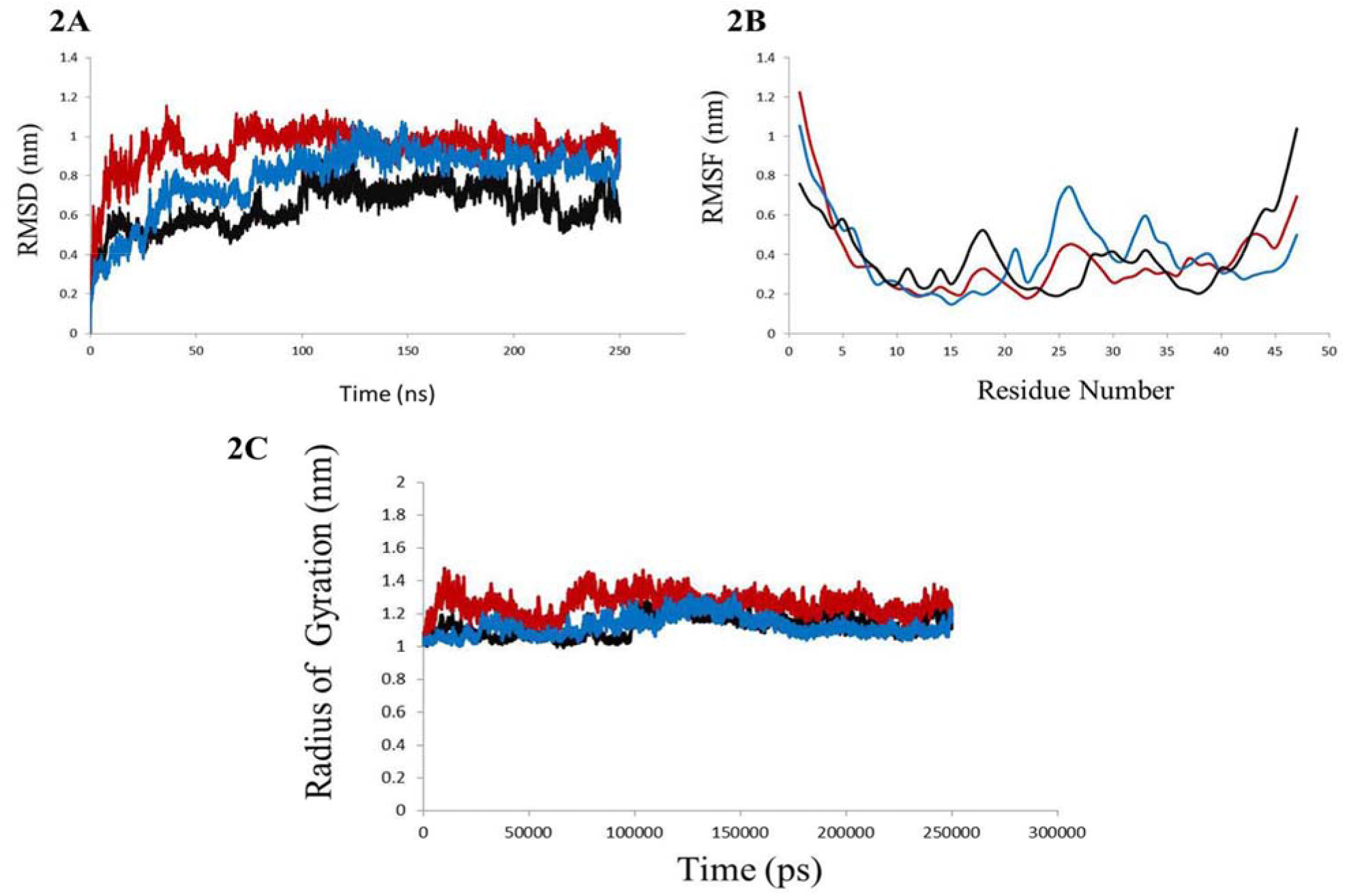
Details of the MD simulation runs in triplicate as inddicated by different colour codes. 2A: Plot of RMSD of the backbone atoms of the peptide during the course of the MD simulations. 2B: Plot of RMSD of the backbone atoms of the peptide during the course of the MD simulations. 2C. Plot of Rg of the peptide during the course of the MD simulations

Residue wise fluctuation is analyzed to check how the amino acid residues would behave during the MD simulation runs. It also measures the local structural flexibility of the peptide. From Fig 2B, it is apparent that the fluctuations of the amino acid residues at both the termini of the peptide were higher. Additionally, the region spanning the amino acid residues 25 to 30 would show higher orders of flexibility.

Fig2C is the plot of Rg with the simulation time. The Radius of gyration is a measure of the distance between the center of mass of the peptide to both the termini. In other words, it would depict the compactness of the peptide. The peptide would attain a compact structure after 100ns (100000ps) of the MD simulation runs.

To further investigate the structural fluidity of the peptide, we tried to analyze the evolution of the secondary structural elements of the peptide during the course of the MD simulation runs. For that we used the tool DSSP. The analysis of the evolution of the secondary structural elements is presented in supplementary figure S3. It was observed that the α-helix at the N-terminal end of the peptide, spanning the amino acid residues 10 to 17, is stably maintained throughout the course of the MD simulations. Similarly, the sheet region spanning the amino acid residues 25 to 30 is a stable secondary structure of the peptide.

### 3.3. Prediction of the ligand binding site on the peptide

The peptide is known to interact with the protein ARNT to exhibit its anti-angiogenic activity [16]. However, we wanted to check whether the peptide would be able to bind to some other ligands as well. Therefore, we tried to extract the binding site information of the peptide using four different tools (Table1). Binding site information is essential for a better understanding of the peptide-drug interactions.

From the table1, it can be safely concluded that the peptide has some additional binding regions centered mostly around its N-terminal part. The amino acid residues at the N-terminal helix have their tendencies to bind charged ligands (table1). The helices are known to have defined polarities which would make this region fit to bind charged ligands.

**Table 1:**
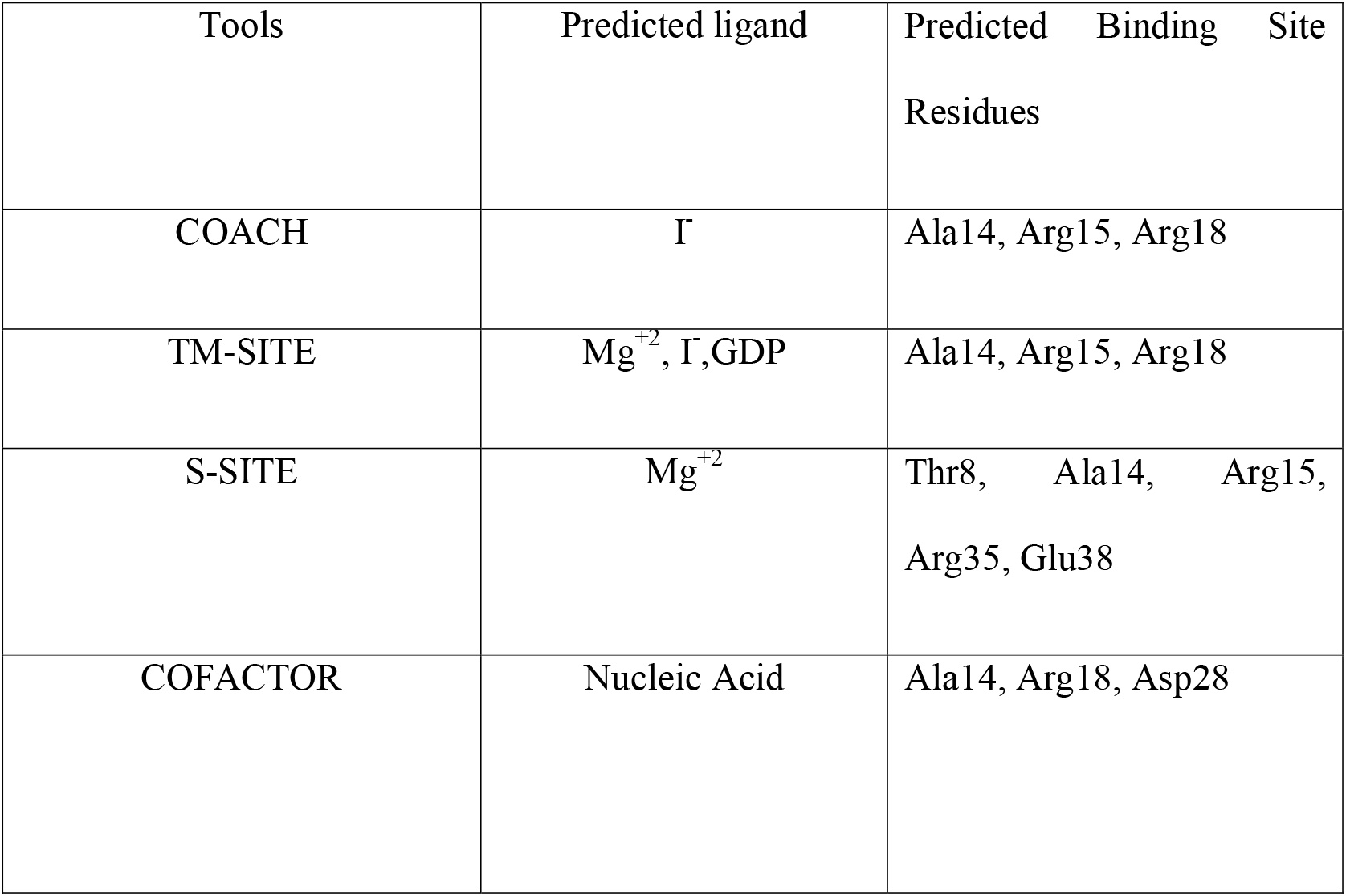
List of the predicted amino acid residues present at the binding site of the peptide and the plausible ligands.

## 4. Discussion

The circular RNAs are associated with a number of different cellular activities. However, circular RNAs belong to the class of non-coding RNAs [1–10]. Surprisingly, the particular circular RNA, circ-0000437, has protein coding potentials and it codes for the peptide CORO1C-47aa. The peptide has its role in prevention of endometrial tumor by binding to ARNT protein [16]. However, till date only the sequence information of the peptide is available. Therefore, we tried to utilize the techniques of bioinformatics to structurally characterize the peptide. Due to the un-availability of suitable templates, the structure of the peptide was built by threading methods. We used MD simulations in triplicate to determine how the peptide would behave in a virtually created cellular environment. We could predict that the peptide is a stable one and mostly hydrophilic. There are stable helical regions at the N-terminal region of the peptide. The central part of the peptide has a stable disposition of sheet regions. The peptide structure did not lose its integrity as observed from the analysis of the Rg values. This would further establish the fact that the peptide would follow a stable folding pattern. We also analyzed the peptide structure to detect the presence of some additional binding sites. We observed that the peptide can bind some charged ligands at its N-terminal part. All these information have been predicted for the first time from our analysis. Since, this peptide has a profound biological significance, our study would be able to shed light on the molecular mechanisms behind the cellular processes where this peptide participates. On top of that, the peptide has anti-tumor activity during the onset of endometrial cancer tumors. This structure-function relationship analysis would be beneficial in predicting the natures of the drugs for the treatment of endometrial cancer tumors.

## 6. Acknowledgement

The authors acknowledge the infrastructural support from the DBT funded BIF centre, University of Kalyani, DST-PURSE, Department of Science & Technology and UGC-SAP, Govt. of India. The author, S.B. is thankful to University of Kalyani for the fellowship provided (URS–SRF). G.B. is thankful to DBT for the BINC fellowship.

## 8. Statements and Declarations

### 8.3. Author contributions

Conceptualization by A.B, methodology by S.B, A.B, formal analysis & investigation by S.B, G.B, writing, review & editing by A.B, G.B. and S.B funding by KU, DST PURSE, SAP-UGC. Resources BIF centre KU, Supervision by A.B.

## Figure legends

**Supplementary figures S1.**
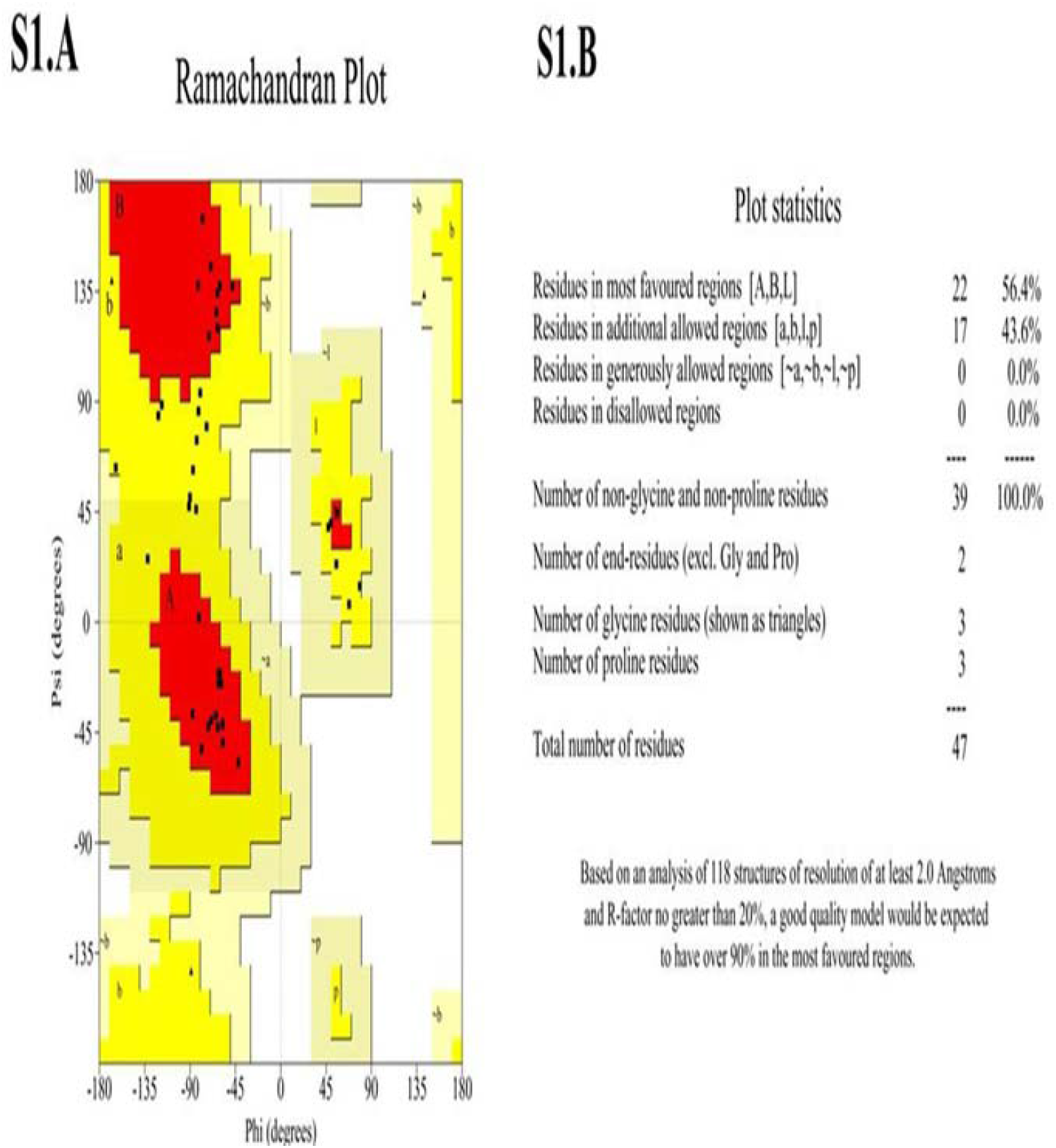
A and S1.B: The Ramachandrand plot of the modeled peptide and the plot statistics, respectively.

**Supplementary figures S2.**
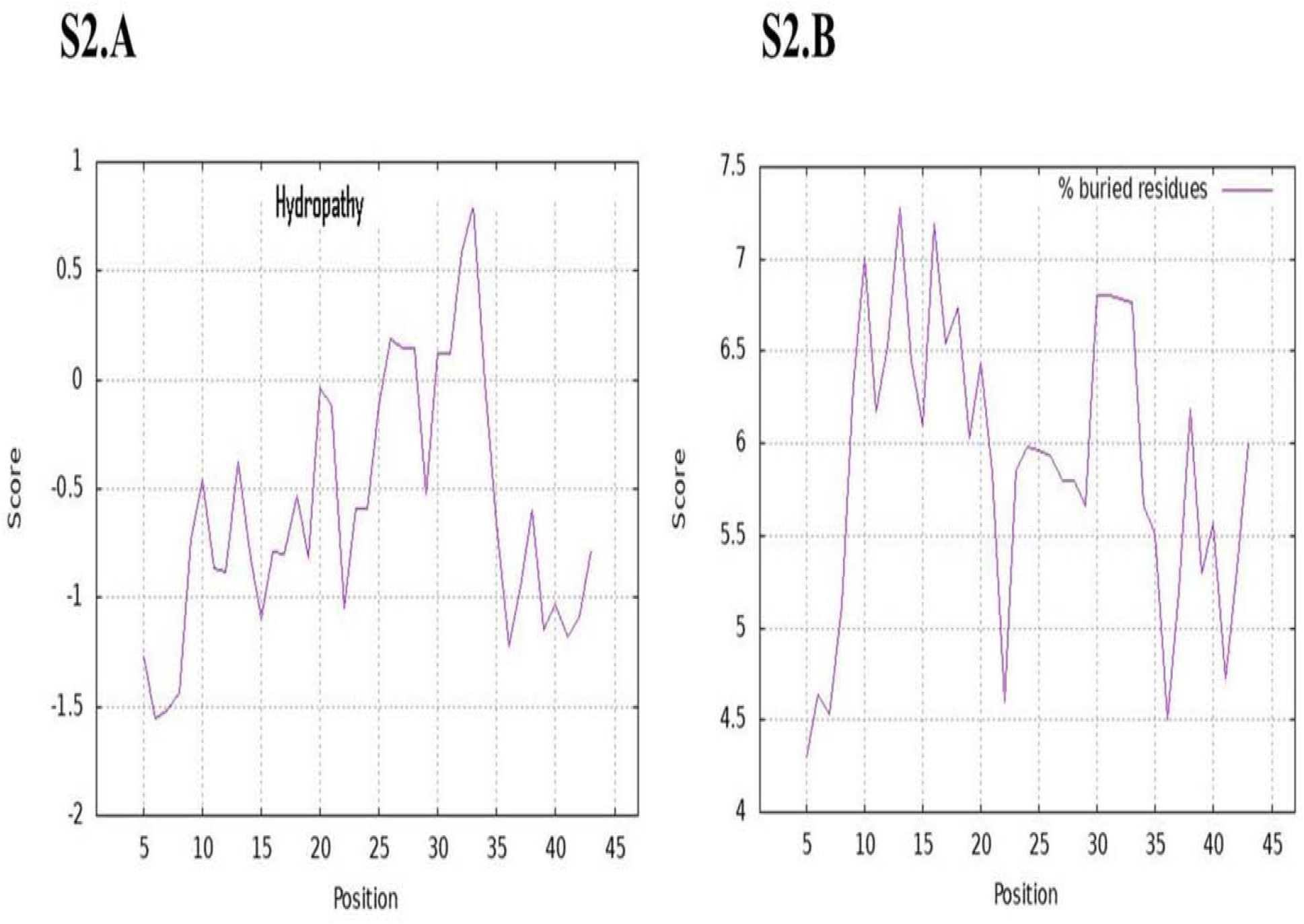
A & S2.B: The relative hydropathy of the amino acids (S2.A) and the percentage of buried residues (S2.B) in the peptide.

**Supplementary figure S3.**
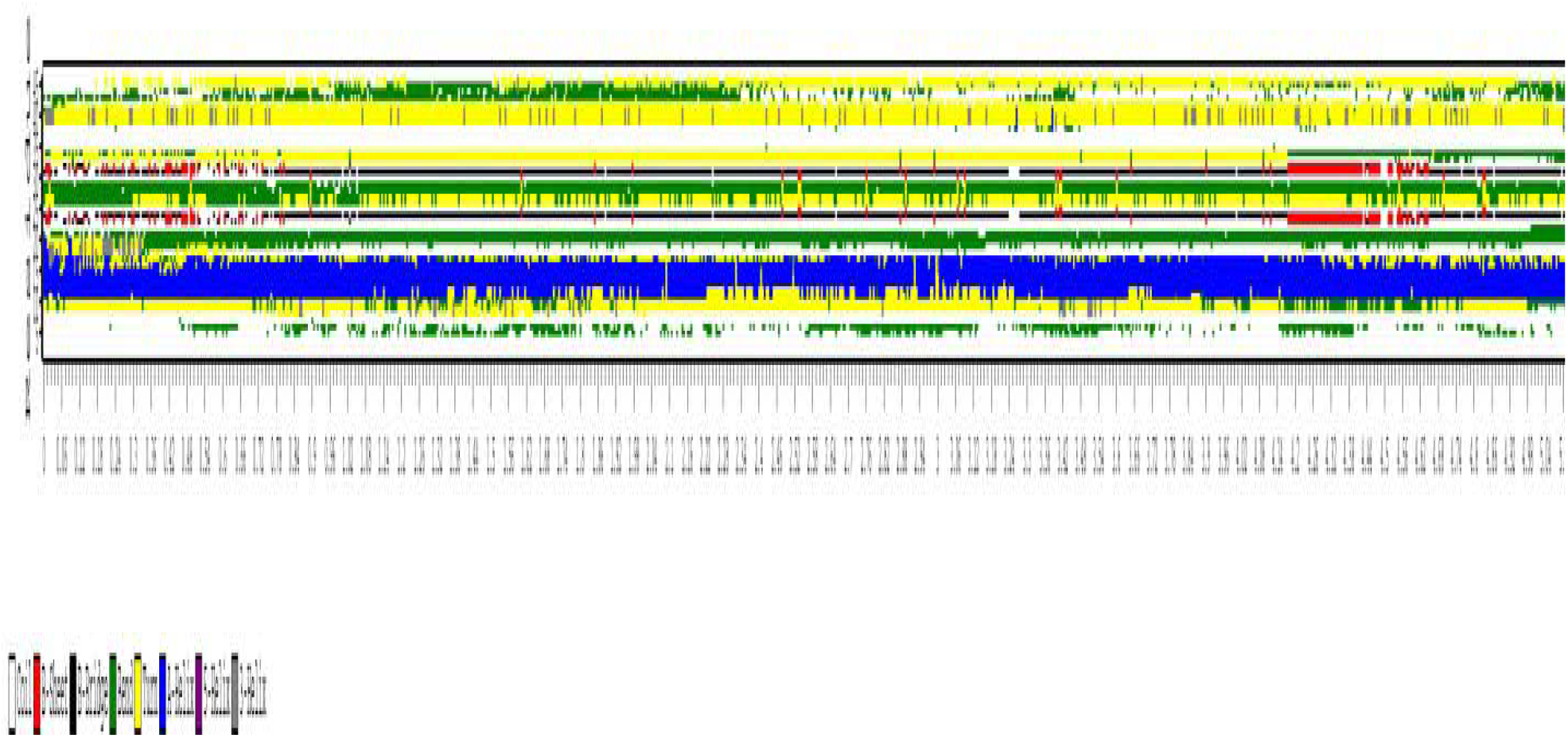
Evolution of the secondary structural elements during the course of MD simulation runs

## Table legend

**Supplementary Table S1.** Sequence analysis of the peptide

| **Parameters** | **Results observed** |
| --- | --- |
| Molecular weight | 5154.80 Da |
| p.I | 9.49 |
| Estimated half life | 30 hours |
| The instability index | 28.83 |
| Aliphatic index | 51.91 |
| Grand average of hydropathicity (GRAVY) | -0.653 |

